# Conscious perception modulates perceptual echoes

**DOI:** 10.1101/2020.08.06.239202

**Authors:** Canhuang Luo, Rufin VanRullen, Andrea Alamia

## Abstract

Alpha rhythms (~10Hz) in the human brain are classically associated with idling activities, being predominantly observed during quiet restfulness with closed eyes. However, recent studies demonstrated that alpha (~10Hz) rhythms can directly relate to visual stimulation, resulting in oscillations which can last for as long as one second. This alpha reverberation, dubbed Perceptual Echoes (PE), suggests that the visual system actively samples and processes visual information within the alpha-band frequency. Although PE have been linked to various visual functions, their underlying mechanisms and functional role are not completely understood. In the current study, we investigated whether conscious perception modulates the generation and the amplitude of PE. Specifically, we displayed two colored Gabor patches with different orientations on opposite sides of the screen, and using a set of dichoptic mirrors we induced a binocular rivalry between the two stimuli. We asked participants to continuously report which one of two Gabor patches they consciously perceived, while recording their EEG signals. Importantly, the luminance of each patch fluctuated randomly over time, generating random sequences from which we estimated two impulse-response functions (IRFs) reflecting the perceptual echoes generated by the perceived (dominant) and non-perceived (suppressed) stimulus respectively. We found that the alpha power of the PE generated by the consciously perceived stimulus was comparable with that of the PE generated during monocular vision (control condition), and significantly higher than the PE induced by the suppressed stimulus. Moreover, confirming previous findings, we found that all PEs propagated as a travelling wave from posterior to frontal brain regions, irrespective of conscious perception. All in all our results demonstrate that conscious perception modulates PE, suggesting that the synchronization of neural activity plays an important role in visual sampling and conscious perception.

## Introduction

The alpha rhythms [8-12 Hz] is the most prominent oscillation in the human brain, and the first one to be described in human electrophysiological recordings (Berger, 1933). It involves most of the cortical regions, but it is most dominant in occipital and parietal areas. Its origin can be related to different processes: some studies pointed at the closed-loop interaction between cortical and thalamic regions, the latter acting as alpha pacemakers (Bollimunta et al., 2011; Lopes da Silva et al., 1980, 1973), but recent evidence indicated uniquely cortical mechanisms as responsible for its generation (Halgren et al., 2019). Just like distinct sources can produce alpha-band rhythms, similarly these alpha band oscillations are likely to serve different functions. On the one hand, alpha oscillations have been shown to strongly but negatively correlate with task demand and increasing attention, hence their presumed involvement in inhibitory functions (Gazzaley and Nobre, 2012; Jensen and Mazaheri, 2010; Klimesch, 2012). On the other hand, alpha waves have been related to information processing, such as the temporal parsing of sensory information (Klimesch et al., 2007) or the perception of visual stimuli (VanRullen, 2016). Regarding the latter, electrophysiological recordings demonstrate that visual stimuli reverberate in visual cortical areas around 10Hz, producing what has been dubbed as perceptual echoes (VanRullen, 2016; Vanrullen and MacDonald, 2012).

Perceptual echoes (PE) are best observed by cross-correlating a non-periodic flickering stimulus, for example a disk whose luminance randomly varies over time, with the EEG signals recorded in occipital and parietal regions. The cross-correlation provides an Impulse Response Function (IRF) which describes the brain response to each stimulus transient. Such response reveals a clear oscillation in the alpha-band whose duration can last for as long as one second. A recent study ascribed the mechanisms generating the echoes to the interactions between brain regions within a predictive coding framework (Alamia and VanRullen, 2019). However, whether PE are a by-product of cortical interactions or serve some specific cognitive function remains unclear. Experimental studies demonstrated that PE are enhanced when repetitions are embedded in the visual sequence, suggesting that they could reflect a role in regularity learning (Chang et al., 2017), whereas other evidence show that attended stimuli generate larger echoes than unattended ones, suggesting that attention allocation plays a role in modulating PE amplitude (Vanrullen and MacDonald, 2012). In addition, PE have been characterized as travelling waves that propagates from occipital to frontal regions, thus including a spatial component that may reflect the hierarchical processing of visual information along the visual system (Alamia and VanRullen, 2019; Lozano-Soldevilla and VanRullen, 2019).

All these findings indicate that PE are relevant to different functional roles in visual information processing, suggesting that they may reflect some fundamental mechanism in cortical processing. In this study we take one step further in this direction by exploring whether PE are modulated by conscious perception. In order to address this question, we tested a pool of participants within a Binocular Rivalry design, in which two different stimuli are shown separately to each eye, generating a rivalry that is resolved with the perception of only one of the two stimuli. In this experiment, a green and a red Gabor patch, respectively tilted by ±45°, were displayed on the left and right side of the screen. We employed a dichoptic mirrors setup to project each stimulus separately to each eye. Importantly, the luminance of each stimulus varied over time in a random, non-periodic way, generating two flickering luminance sequences. Participants were instructed to continuously report which colored Gabor patch was being perceived throughout the experiment, thus defining two sequences corresponding to the dominant and suppressed stimuli. Here, we aimed at computing the echoes by cross-correlating the EEG recordings with each sequence, in order to assess whether the generation and amplitude of PE is modulated by conscious perception.

## Methods

### Participants and statistical power analysis

We estimated the number of participants via a statistical power analysis based on previously published data investigating PE in binocular vision (Brüers and VanRullen, 2017). We determined the effect size as equal to 1.7, computed as the mean alpha power difference between the actual echoes and the ones obtained after shuffling the temporal sequences (i.e. surrogate echoes, see below). Setting the power level to 0.90 and the statistical threshold at 0.05, that effect size requires a number of participants equal to 4. However, considering that we based our effect size estimates on binocular vision (i.e. both eyes fully perceiving the same stimulus), and we aimed at computing echoes in condition of monocular vision and binocular rivalry, we tripled the estimate, including a final number of 12 participants (7 female, mean age 26, SE=0.9). All participants had normal or corrected-to-normal vision and gave written consent before the first session of the experiment, in accordance with the Declaration of Helsinki. This study was carried out in accordance with the guidelines for research at the “Centre de Recherche Cerveau et Cognition” and the protocol was approved by the committee “Comité de protection des Personnes Sud Méditerranée 1” (ethics approval number N° 2016-A01937-44).

### Experimental procedure

Each participant completed two sessions on two different days. One session consisted of 10 blocks, each composed of 10 trials. A trial lasted for 30s each. The design consisted of two conditions: in half of the blocks, including the first one, participants performed Binocular Rivalry (BR) trials, whereas on every other block they performed Physical Alternation (PA) ones. In BR trials, two Gabor patches, each encircled by a square frame (visual angle of the patch: 4 degrees, visual angle of the frame: 4.5 degrees), were shown separately to each participant’s eyes. Patches were different in color and inclination, either red or green, ±45°, the color-inclination associations were kept constant throughout each experiment, but randomized between participants. The color and orientation of the stimulus served mainly to help identify the perceived stimulus and thereby facilitate perceptual reports from the participants, however the main experimental variable was the stimulus luminance. The luminance of the Gabor patches changed randomly over time, and this random sequence was designed to have the same spectral power at every frequency (fig.1A). Importantly, the range in the two colors luminance was carefully calibrated and equalized to avoid any perceptual biases. The physical position on the screen of the two Gabor patches was switched on each trial (i.e. either the left or the right side). The task was to report which patch was perceived by moving a joystick either to the left or the right (e.g., one participant instructions were to lean the joystick to the left when green was perceived and to the right when red was perceived). The color-side associations were pseudorandomized between participants. Importantly, participants were encouraged to account continuously for their visual perception, reporting intermediate joystick positions when the perception of both patches overlapped. Each trial started by pressing a joystick button, and participants were encouraged to rest between trials. Each BR block was followed by a PA one. In PA blocks only one Gabor patch was displayed at a time, replaying the exact sequence of Gabor patches reported in the previous BR block. The task’s instructions were the same, with the exception that participants were no longer performing a binocular rivalry task. The goal of such replays was to estimate precisely the reaction time in each trial, and correctly segment the actual perception in the BR blocks before computing the echoes. Moreover, PA blocks served as a control condition to assess PE in condition of monocular vision, without rivalry.

**Figure 1.**
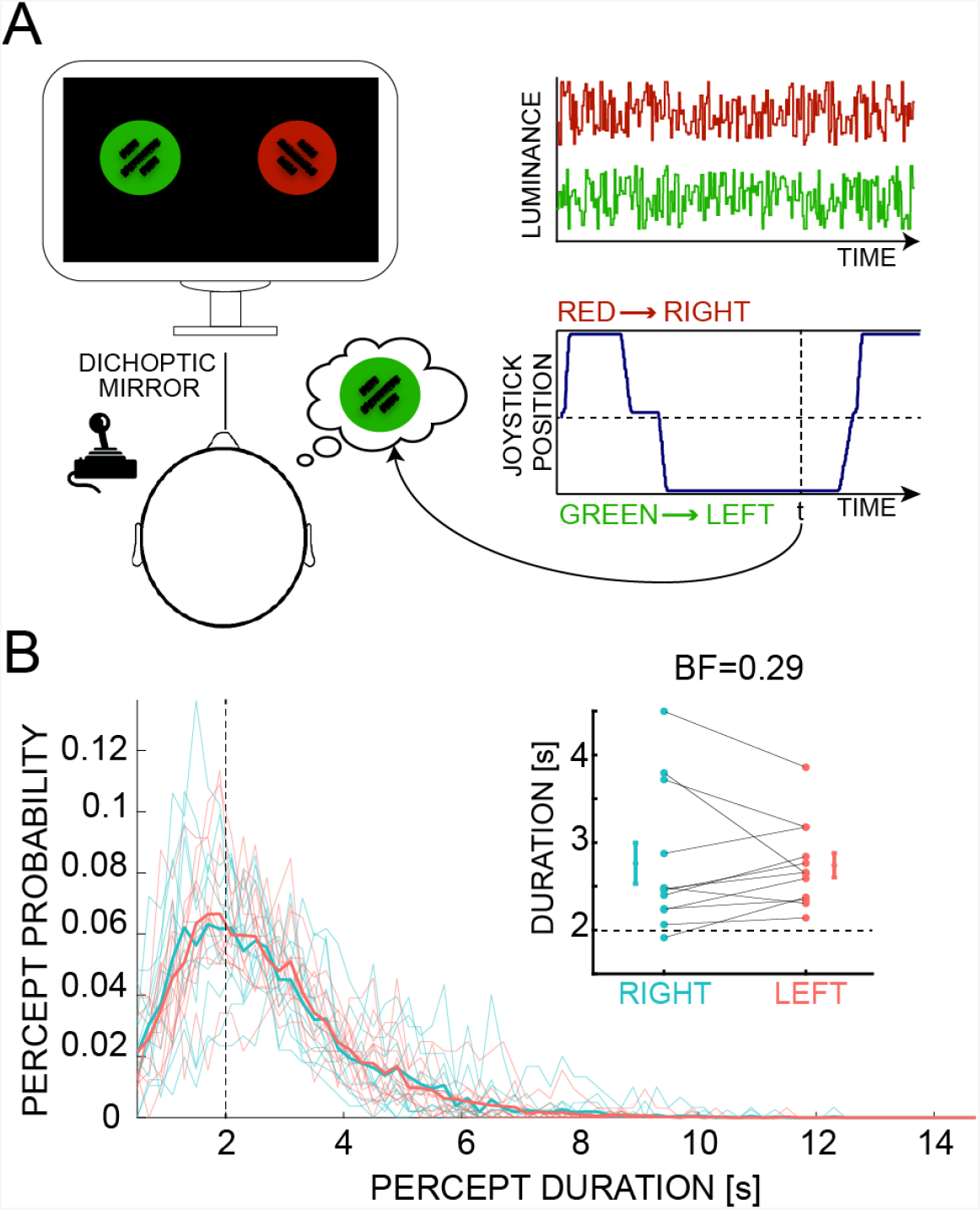
Experimental design. A) Participants stared at the screen through a set of dichoptic mirrors that projected the left and right side of the screen to the left and right eye respectively. Two stimuli, placed on the two sides of the screen, were Gabor patches of different color and orientation, either red or green with a ± 45° angle. Participants reported which patch they perceived by moving a joystick to either side, each one associated to a stimulus (pseudo-randomly between participants, consistent across blocks and sessions). B) Distribution of percept duration in seconds. On average, participants perceived one or the other stimulus for 2 seconds. We discarded percepts below this threshold in all analyses. Overall there was no difference in the duration of stimuli placed to the left and to the right (Bayesian t-test, BF_10_ = 0.29, error=0.021%).

### EEG recording and analysis

#### Recording and Preprocessing

EEG signals were recorded using a 64-channel active BioSemi EEG system (1024Hz sampling rate), and 3 additional ocular electrodes were used. The preprocessing was performed in EEGlab (Delorme and Makeig, 2004) and consisted first in down-sampling the data to 160Hz followed by a high-pass (>1Hz) and a notch (47-53Hz) filter. Data were then average re-referenced and segmented from 200ms before trial beginning until its end (-200ms to 30,000ms). Each epoch was then baseline corrected by subtracting the average between −200ms and stimulus onset.

#### Perceptual Echoes

PE are computed by cross-correlating the luminance sequences with the corresponding EEG signal. As reported in previous studies, PE are stronger in occipital and parietal regions (VanRullen, 2016; Vanrullen and MacDonald, 2012), hence we focused our analysis on signals recorded in POz (note that similar results are obtained when using the electrodes Oz and Pz). First, the reaction time (RT) for each response was estimated from the PA condition, as we knew the exact time when the stimuli were presented to the subjects. We used these RT estimates to infer the exact timing of the perceptual switch during the BR condition, that is we shifted backward the responses in BR to account for the reaction times. Next, we segmented the EEG signals and the corresponding sequences according to participants’ perceptions. In order to identify the temporal segments in which participants reported a full perception (either left or right), we normalized joystick responses between −1 (left) and 1 (right), and we included all the sequences in which the response was above a threshold set to ±0.95 and longer than 2 seconds to ensure the sequences were long enough for the reliable estimation of PE (figure 1B). In BR blocks, for each segment we cross-correlated the EEG signal with the sequence of the perceived patch (i.e. dominant) and the non-perceived patch (i.e. suppressed). In PA blocks, we cross-correlated the EEG signal with the one sequence shown. In both conditions, the cross-correlation was computed on lags between −0.5 and 2 seconds. The module of the PE spectra was computed with a Fast Fourier transform over the delays between 0.25 and 1 second. From each spectra we extracted the average power in the alpha-band [8-12Hz]. To estimate a baseline for comparison, we computed the same power spectra in surrogate echoes, obtained by cross-correlating the EEG signals with the luminance sequences after having shuffled their temporal order. Lastly, we compared the amount of alpha power in the echoes between conditions (dominant, suppressed and physical alternation) by means of a Bayesian ANOVA having CONDITION as a fixed factor and subject as a random term. For dependent variable we considered the amount of alpha power computed in decibel [dB] as:

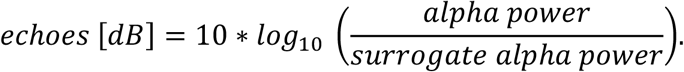

Regarding the time-frequency analysis we computed the power-spectra using a wavelet transformation (1-40 Hz, in log-space frequency steps with 1-20 cycles) of each IRF (i.e. the result of cross-correlating each luminance sequence and the temporally-aligned EEG signal). We applied a baseline correction by subtracting the average activity 200 ms prior to 0 lags, and we extracted the mean value in the alpha range [8-12Hz] in the time-window between 250ms and 850ms. As previously, we computed a Bayesian ANOVA to compare the alpha power spectra between each condition.

#### Travelling waves analysis

We eventually assessed how PE propagate through cortex as a travelling wave. As in our previous studies (Alamia and VanRullen, 2019; Pang et al., 2020) we computed the echoes in seven midline electrodes (Oz, POz, Pz, CPz, Cz, FCz, Fz), and we created 2D maps by stacking signals from those electrodes (see figure 4A). From the 2D map we computed a 2D-FFT, in which the power of the upper left quadrant represents the amount of waves travelling in a forward direction (FW - from occipital to frontal electrodes) whereas the lower left quadrant quantifies the amount of waves travelling backward (BW - from frontal to occipital). Note that the same values can be found in the right quadrants, since the 2D-FFT is symmetric around the origin. Since we investigated the propagation of the alpha-band echoes, we extracted the maximum values within the alpha range [8-12Hz]. In order to quantify the amount of waves above chance level, we computed a surrogate distribution of values by shuffling the electrodes order before quantifying the 2D-FFT (obtaining FW_ss_ and BW_ss_ for forward and backward waves respectively). Similarly to the previous analyses, we computed the amount of waves in decibel [dB] for FW and BW waves according to the following formula:

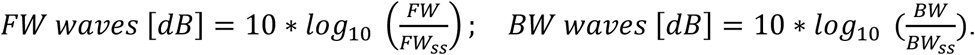

#### Statistical analyses

All statistical tests were performed within the Bayesian framework, assessing the likelihood of a model given the data. This analysis produces a Bayes Factor (BF), which quantifies the ratio between models testing the alternative over the null hypothesis. Throughout the paper, all BFs comply with this convention – i.e. the probability of the alternative hypothesis over the null hypothesis, usually indicated as BF_10_. In practice, a large BF (~BF>3) provides evidence in favor of the alternative hypothesis, whereas low BF (~BF<0.3) suggests a lack of effect (Masson, 2011; Smith, 2001). All analyses were performed in JASP (JASP Team, 2018; Love et al., 2015).

## Results

### Echoes

The main goal of this study was to determine whether PE are influenced by conscious perception. First, we estimated the averaged alpha-band power of the PE generated by the dominant and suppressed stimuli, along with those measured during the physical alternation (PA) task. Each PE was obtained by cross-correlating the EEG recording (POz electrode) with the corresponding luminance sequence (Fig 2). In order to quantify the power in the alpha range in each condition (figure 3A) we computed the corresponding surrogate values after shuffling the temporal order of the sequence, thus expressing the PE alpha amplitude as a ratio measured in dB (see Methods for details). The graph in figure 3B reveals a significant difference between conditions, as confirmed by a Bayesian ANOVA (CONDITION factor: BF_10_ = 9.442, error=0.411%). A post-hoc Bayesian t-test comparison confirms a significant difference between dominant and suppressed echoes (BF_10_ = 23.926, error < 0.001%). Moreover, echoes generated in the PA conditions were larger than the one generated by the suppressed sequence (BF_10_ = 3.276, error < 0.001%), but we observed no difference between PE generated by the dominant sequence and in the PA conditions (BF_10_ = 0.159, error < 0.001%). Interestingly though, both dominant and suppressed echoes proved to be significantly larger than zero (Bayesian one sample t-test, both dominant and suppressed BF_10_ >> 100, error < 0.001%), suggesting that PE can be elicited also without conscious perception. Not surprisingly, we also found significant echoes in the Physical Alternation conditions, i.e. when only one Gabor was displayed (BF_10_ >> 100, error < 0.001%), confirming the results of a recent study (Schwenk et al., 2020) showing that PE, although strongly reduced, can still be observed in conditions of monocular vision.

**Figure 2.**
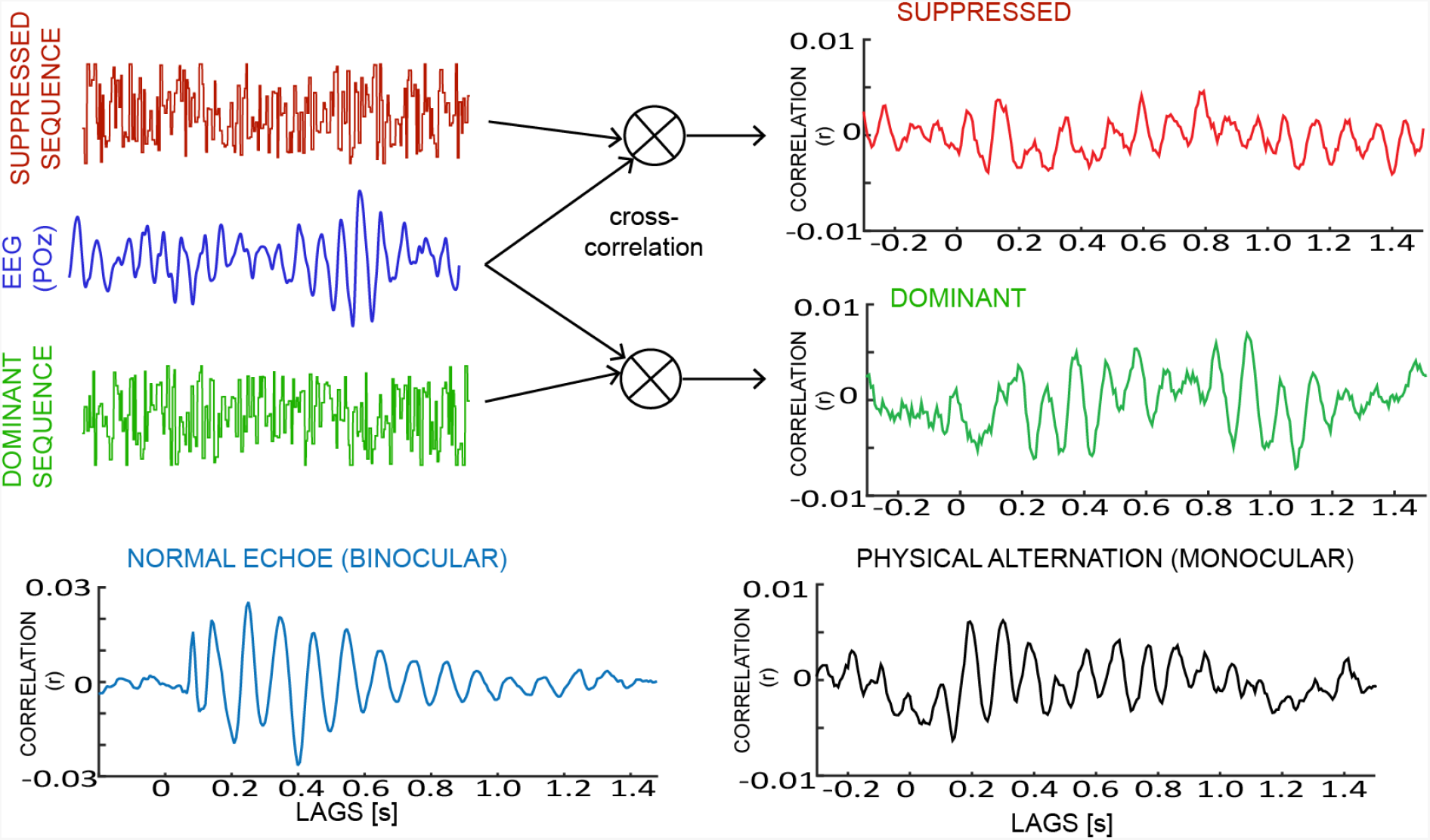
Computing perceptual echoes. Two random (independent) temporal sequences of luminance were displayed on opposite sides of the screen. Given the dichoptic mirror setup, each sequence was perceived by one eye only, producing a binocular rivalry that was resolved with one perceived sequence (i.e. dominant, in green) and one non-perceived (i.e. suppressed, in red). We computed PE by cross-correlating each sequence and the corresponding EEG signal (POz electrode), revealing a reverberation in the alpha-band interval. The same procedure was used to compute the PE in the Physical Alternation condition (in black). The bottom-left panel shows for comparison a PE computed in case of binocular vision (Brüers and VanRullen, 2018). Note the difference in the y-axis.

**Figure 3.**
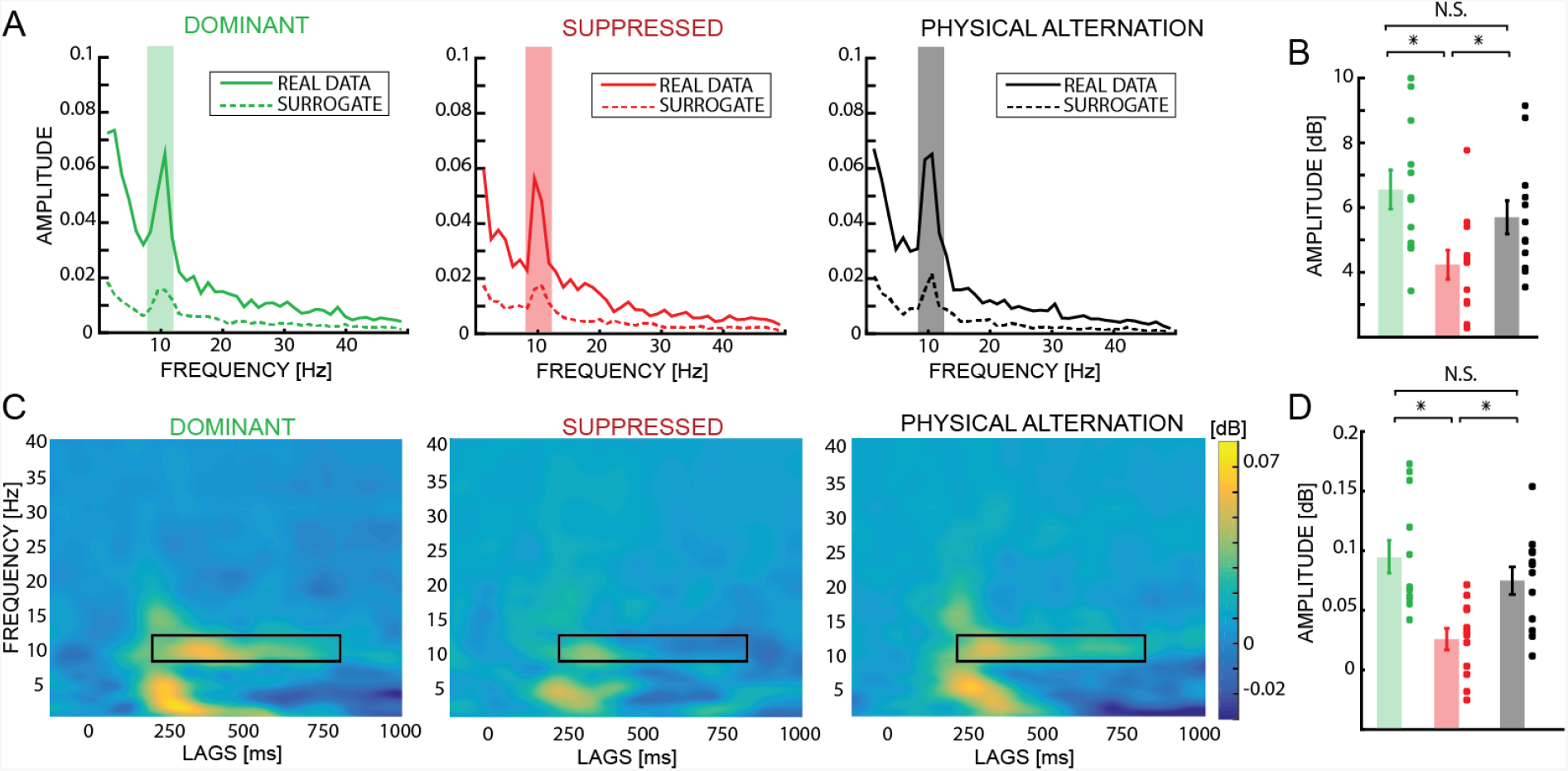
Echoes and time-frequency results. A) The figure shows the power spectra of the PE obtained in the dominant (green), suppressed (red) and PA (black) condition. The dashed line is the spectra obtained in the surrogate echoes, i.e. computed after having shuffle the temporal order of the luminance sequence. We focused the analysis in the alpha range (shaded in each panel). B) The difference in the power spectra in the alpha range expressed in dB. The plot reveals a significant difference between suppressed and both dominant and PA conditions, revealing that conscious perception increases the amplitude of the PE. However, all the conditions have PE larger than chance. C) Time-frequency spectrogram for each condition (color-code as in A). Dominant and PA show a larger amplitude than in suppressed, as confirmed by the plot in D) comparing the amplitude in the time-frequency region of interest (black box) in each condition.

### Time-Frequency

In order to assess the temporal dynamics of the PE in each condition (i.e. dominant, suppressed and physical alternation) we performed a time-frequency analysis on the echoes. In line with our previous results, figure 3C reveals a stronger effect in the alpha band in the dominant and PA conditions compared to the suppressed condition. We confirmed this result by computing per each subject the average power within the temporal window [250ms – 850ms], in the alpha range [8Hz – 12Hz]. As in the previous analysis, a Bayesian ANOVA confirmed a significant difference between the three conditions (CONDITION factor: BF_10_ = 54.22, error=0.011%), as well as the post-hoc Bayesian t-test which positively confirms a difference between dominant and suppressed (BF_10_ >> 100, error < 0.001%) and PA and suppressed (BF_10_ = 13.15, error < 0.001%), with mild evidence in favor of no difference between dominant and PA (BF_10_ = 0.414, error = 0.022%). Moreover, all the conditions proved significantly larger than zero, even though much larger Bayes Factors were observed in the dominant and PA conditions (dominant and PA: BF_10_ >>100, error < 0.001%; suppressed BF_10_ = 5.769, error < 0.001%). Overall the time-frequency analysis confirmed the previous results, indicating that PE elicited in the dominant and PA conditions are larger than in the suppressed condition.

### Travelling waves

Eventually we investigated whether PE elicited during binocular rivalry propagate through cortex as forward travelling waves (i.e. from occipital to frontal regions), as recently showed in the case of binocular vision (Alamia and VanRullen, 2019; Lozano-Soldevilla and VanRullen, 2019). We quantified the amount of forward and backward waves as shown in figure 4 obtaining for each participant a value in dB for each condition (see methods for details). Interestingly, a Bayesian ANOVA performed with factors DIRECTION (FW and BW) and CONDITION (dominant, suppressed and PA) revealed a significant difference between FW and BW waves (BF_10_ = 3.963, error = 0.001%) but neither a difference between conditions (BF_10_ = 0.100, error = 0.007%), nor an interaction (BF_10_ = 0.097, error = 0.012%). A Bayesian t-test comparing the amount of FW waves against zero confirmed that PE propagates from occipital to frontal regions when elicit by dominant (BF_10_ = 6.783, error < 0.001%), suppressed (BF_10_ = 18.907, error < 0.001%) and monocular sequences (PA, BF_10_ = 3.655, error = 0.001%).

**Figure 4.**
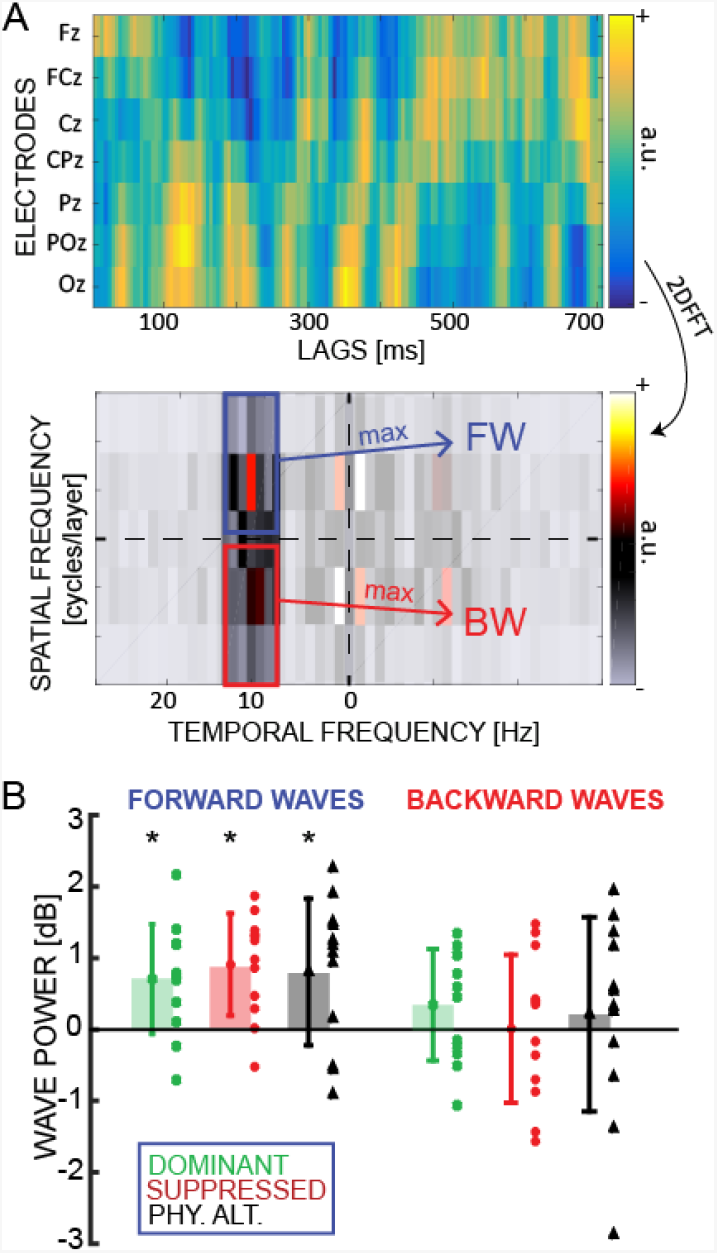
Travelling waves results. A) We first obtain 2D maps stacking the PEs recorded over the 7 midline electrodes. The color code indicates the PE amplitude. From the 2DFFT we computed the amount of FW and BW waves in each condition as the maximum amplitude value in the corresponding quadrant (restricted to alpha-band frequencies). B) The results –expressed in dB, i.e. corrected using the surrogate, see methods-show that PE travel as FW waves in all conditions, irrespective of the conscious perception of the stimulus.

## Discussion

Previous studies showed that visual information reverberates in posterior brain regions in the alpha-band frequency range, as observed by cross-correlating white noise luminance sequence with EEG recordings (VanRullen, 2016; Vanrullen and MacDonald, 2012). Such reverberation, dubbed as Perceptual Echoes (PE), proved in several studies to be related to various cognitive functions, such as attention (Vanrullen and MacDonald, 2012) and statistical learning (Chang et al., 2017). In this study we investigated whether PE can be modulated by conscious visual perception using a binocular rivalry design. Our results indicate that PE can be generated by both consciously perceived and suppressed stimuli, but the former elicit larger PE than the latter, and of comparable amplitude as the PE generated during monocular vision. Moreover, we reported that the PE generated by both conscious and unconscious visual perception propagates as a travelling wave from occipital to frontal regions, possibly reflecting bottom-up processing in the visual system (Alamia and VanRullen, 2019; Lozano-Soldevilla and VanRullen, 2019; Pang et al., 2020).

Similar to the finding that PEs are enhanced by attention (Vanrullen and MacDonald, 2012), we found that PEs generated by the consciously perceived sequence (i.e. the dominant stimulus) contain larger alpha power. If on the one hand previous studies demonstrated that binocular rivalry might be modulated by attentional mechanisms (Zhang et al., 2011), on the other hand the alpha power enhancement we observed in the PE in the dominant condition is unlikely to be due to selective attentional processes, as participants were not instructed to pay attention specifically to one of the two stimuli. Yet, we can de facto assume that the attentional focus is driven to the stimulus perceived as conscious, thus increasing the PE alpha power as compared to the suppressed one. Interestingly, previous studies reported opposite effects of attention on alpha power: while attention decreases stimulus non-specific alpha power, it nonetheless increases the spectral (alpha-band) power of PE (Thut et al., 2006; Vanrullen and MacDonald, 2012; Worden et al., 2000). It could be interesting then to test the hypothesis that conscious perception plays a similar role in the modulation of alpha power. One possible experimental approach would be to lateralize the suppressed and dominant stimulus, thus assessing whether conscious perception modulates the alpha power in each occipital hemisphere similarly to attention. Further experiments will shed light on this interesting hypothesis.

The enhancement of PE alpha power in the dominant condition together with the previous finding that attention enhances PE are reminiscent of the application of frequency-tagging in binocular rivalry. Previous studies revealed that both conscious perception and attention allocation increase the spectral power corresponding to the steady state visually evoked potential (SSVEP) (Ding et al., 2006; Srinivasan et al., 1999; Tononi et al., 1998). Even though SSVEP showed similar effects as PE in binocular rivalry and attention tasks, their underlying mechanisms are likely different. SSVEP is a passive brain response to a rhythmic stimulation, reflecting the spectral characteristics of the generating stimulus, whereas PEs are characterized by a clear 10 Hz oscillation without a corresponding 10Hz peak in the visual stimulus, possibly reflecting computational cortical mechanisms (Alamia and VanRullen, 2019). Despite the functional differences, it is tempting to speculate that in both SSVEP and PE, conscious perception modulates the amount of synchronized activity in brain regions: the higher alpha power in dominant PE might be associated with a larger synchronization of local neural activity, which might be akin to the increase in the power of the SSVEP related to conscious perception.

Besides their oscillatory temporal dynamics, PE elicited in conditions of binocular vision have been described in view of their spatial component, characterized as a travelling waves propagating from occipital to frontal regions (Alamia and VanRullen, 2019; Lozano-Soldevilla and VanRullen, 2019). In this study we replicate a similar pattern of results, as we observed the same amount of forward travelling waves in both dominant and suppressed conditions, as well as during the physical alternation task (i.e. monocular vision). Surprisingly, the difference in PE amplitude observed between dominant and suppressed conditions was not reflected in the waves’ directional power, as waves seem to propagate from lower to higher brain regions with the same strength irrespective of conscious perception. Possibly, the relatively poor spatial resolution of EEG recordings prevents us from accurately comparing the two conditions, and different experimental techniques will be required to reveal different directional strengths in the propagation of dominant and suppressed travelling waves. However, one could speculate that PE generated by the consciously perceived sequence (i.e. dominant) propagate further in the visual hierarchy, reaching frontal regions which are supposedly involved in conscious perception (Koch et al., 2016; Miller, 2011), whereas the oscillatory waves of PE generated by the suppressed sequence vanish at an earlier stage of visual processing. Further studies will be needed to fully address this hypothesis.

Several previous studies investigated conscious perception in light of the Predictive Coding framework (Hohwy et al., 2008; Lamme, 2015; Seth et al., 2012; Weilnhammer et al., 2017). Predictive Coding (PC) is an influential scheme in cognitive neuroscience that describes the brain as a hierarchical system, in which higher regions generate predictions about the activity of lower ones, and the difference between predictions and actual activities (i.e. the prediction error) is used to update the upcoming predictions (Huang and Rao, 2011). Is it possible to combine within the same framework predictive coding, conscious perception and PE? In a previous study we demonstrated that a simple model simulating the interaction between brain regions and based on PC principles can account for the generation and propagation of PE as travelling waves (Alamia and VanRullen, 2019), under the assumption of plausible biological constraints (i.e. communication delays and time constants). Interestingly, additional experimental evidence supporting the tie between conscious perception, predictive coding and travelling waves was recently reported in another study investigating how psychedelic drugs altered travelling waves, supposedly by relaxing the weighting of top-down predictions, thereby releasing the bottom-up flow of information carried by sensory input (Alamia et al., 2020). On the other hand, other studies have characterized conscious perception within a PC framework as the consequence of prediction-error minimization (Friston, 2013; Hohwy, 2012; Hohwy et al., 2008; Strauss et al., 2015): this compelling hypothesis advocates that predictions are generated to efficiently explain and interpret the causes underlying our sensory information, thus generating our conscious perception of the world (Panichello et al., 2013). All in all, the result that PE are influenced by conscious perception –as we demonstrated in this study-,leads to the compelling speculation that Predictive Coding is pivotal in the generation of both PE and conscious experiences.

In conclusion, the current study investigated PE by employing binocular rivalry, and revealed that these are modulated by conscious perception, but consciousness is not necessary to elicit them. In addition, PE evoked by both consciously and unconsciously perceived stimuli propagate from occipital to frontal regions as a travelling wave, irrespective of the conscious modulation.

